# Nr4a2 blocks oAβ-mediated synaptic plasticity dysfunction and ameliorates spatial memory deficits in the APP_Sw,Ind_ mouse

**DOI:** 10.1101/2024.01.24.577010

**Authors:** Judit Català-Solsona, Stefano Lutzu, Pablo J. Lituma, Cristina Fábregas-Ordoñez, Dolores Siedlecki, Lydia Giménez-Llort, Alfredo J. Miñano-Molina, Carlos A. Saura, Pablo E. Castillo, José Rodriguez-Álvarez

**Affiliations:** Institut de Neurociències and Dpt. Bioquímica i Biología Molecular, Universitat Autònoma de Barcelona, 08193 Bellaterra, Barcelona, Spain; Centro de Investigación Biomédica en Red sobre Enfermedades Neurodegenerativas (CIBERNED), 28031 Madrid, Spain; Dominick P. Purpura Department of Neuroscience, Albert Einstein College of Medicine, New York, NY 10461, USA; Institut de Neurociències and Dpt. Psiquiatria i Medicina Legal, Universitat Autònoma de Barcelona; Department of Psychiatry & Behavioral Sciences, Albert Einstein College of Medicine, New York, NY 10461, USA

## Abstract

Alzheimer’s disease AD is associated with disruptions in neuronal communication, especially in brain regions crucial for learning and memory, such as the hippocampus. The amyloid hypothesis suggests that the accumulation of amyloid-beta oligomers (oAβ) contributes to synaptic dysfunction by internalisation of synaptic AMPA receptors. Recently, it has been reported that Nr4a2, a member of the Nr4a family of orphan nuclear receptors, plays a role in hippocampal synaptic plasticity by regulating BDNF and synaptic AMPA receptors. Here, we demonstrate that oAβ inhibits activity-dependent Nr4a2 activation in hippocampal neurons, indicating a potential link between oAβ and Nr4a2 down-regulation. Furthermore, we have observed a reduction in Nr4a2 protein levels in postmortem hippocampal tissue samples from early AD stages. Pharmacological activation of Nr4a2 proves effective in preventing oAβ-mediated synaptic depression in the hippocampus. Notably, Nr4a2 overexpression in the hippocampus of AD mouse models ameliorates spatial learning and memory deficits. In conclusion, the findings suggest that oAβ may contribute to early cognitive impairment in AD by blocking Nr4a2 activation, leading to synaptic dysfunction. Thus, our results further support that Nr4a2 activation is a potential therapeutic target to mitigate oAβ-induced synaptic and cognitive impairments in the early stages of Alzheimer’s disease.

## INTRODUCTION

Alzheimer’s disease (AD) is the major cause of dementia in the elderly leading to memory loss, cognitive decline, psychiatric deficits, and neuronal death. Multiple pieces of evidence indicate that cognitive impairment observed in the early stages of AD could be due to a disruption of neuronal communication in brain areas that are essential for learning and memory, such as the hippocampus (Selkoe, 2002; Forner et al., 2017). Several transgenic mouse models of AD present impairments of hippocampal synaptic plasticity associated with the progression of the pathological hallmarks (Chapman et al., 1999; Oddo et al., 2003; Gong et al., 2004; Balducci et al., 2011; Crouzin et al., 2013). The amyloid hypothesis has proposed that naturally secreted Aβ oligomers (oAβ) accumulation would affect synaptic efficacy leading to neuronal loss and dementia (Hardy and Selkoe, 2002) and several studies have suggested that early oAβ action on hippocampal synaptic plasticity could contribute to cognitive dysfunction in AD. For example, soluble oAβ inhibits long-term potentiation (LTP) (Walsh et al., 2002; Shankar et al., 2008; Jo et al., 2011) and facilitates long-term depression (LTD) in the hippocampus, supporting that early oAβ action on hippocampal synapses contributes to early cognitive dysfunction in AD (Forner et al., 2017).

Excitatory synaptic transmission is tightly regulated by the number of glutamate AMPA receptors (AMPARs) at the synapse. Several studies have shown that oAβ-mediated synaptic depression in the hippocampus is linked to an internalisation of synaptic AMPARs, a blockade of the recruitment of synaptic AMPARs by LTP and an alteration of the postsynaptic density proteome (Hsieh et al., 2006; Minano-Molina et al., 2011; Sanderson et al., 2021). A reduction in GluA1-containing AMPARs in APP_Sw,Ind_ mice was observed before the appearance of classical AD pathological hallmarks (β-amyloid plaques and neurofibrillary tangles) (Minano-Molina et al., 2011). These reduction in AMPARs is correlated with the appearance of oAβ species in the brain parenchyma and early deficits in learning (España et al., 2010; Minano-Molina et al., 2011). Moreover, the learning deficits observed at early stages in the APP_Sw,Ind_ mice correlated with an alteration of a transcriptional program related to synaptic transmission and plasticity that depends on the CREB-regulated transcription coactivator-1 (CRTC1) (España et al., 2010; Parra-Damas et al., 2014). This suggests that mechanisms at the gene-regulatory level could be involved in the dysfunction of glutamatergic synapses at early stages of AD (Saura and Valero, 2011). Among the genes down-regulated in the APP_Sw,Ind_ mice we find the Nr4a family of orphan nuclear receptors (Nr4a1, Nr4a2 and Nr4a3) (España et al., 2010; Parra-Damas et al., 2014), which have been linked to hippocampal plasticity. General genetic blockade of the Nr4a family produced an impairment of hippocampal LTP and severe deficits in hippocampal-dependent contextual fear memory (Hawk et al., 2012; Bridi and Abel, 2013). In particular, Nr4a2 is up-regulated in the hippocampus following learning of spatial tasks (Aldavert-Vera et al., 2013), and its overexpression ameliorates cognitive impairment in aging (Kwapis et al., 2019). The mechanisms involved in Nr4a2-mediated hippocampal synaptic plasticityare poorly understood. Recently, we reported that neuronal activity triggers an ionotropic glutamate receptor/Ca^2+^/CREB-CRTC1 pathway that mediates Nr4a2 activation which modulates hippocampal synaptic plasticity by increasing BDNF and synaptic AMPARs (Català-Solsona et al., 2023).

The reduction of Nr4a2 mRNA in human AD brain samples (Parra-Damas et al., 2014; Moon et al., 2019) and the Nr4a2-mediated regulation of synaptic AMPARs, suggests that AD-associated Nr4a2 down-regulation could contribute to early synaptic dysfunction observed in experimental models of AD. In fact, it has been described that the number of Nr4a2-expressing cells is significantly reduced in the 5XFAD mouse with age while Aβ plaques deposition increases (Moon et al., 2014). Also, a reduction of Nr4a2 expression by shRNA injection in the subiculum of 5XFAD mice, produced an increase in the size of the Aβ plaques, a decrease in Neu-N positive cells and an increase in reactive microglia. Consistent with these findings, overexpression of Nr4a2 or administration of Nr4a2 activators in the subiculum were able to reverse these effects (Moon et al., 2019).

In the present study, we further studied the link between Nr4a2 and AD-associated synaptic and cognitive impairments and whether oAβ could be responsible for the decrease in Nr4a2 expression observed in AD. We found that oAβ blocked activity-dependent Nr4a2 activation in hippocampal neurons and that pharmacological Nr4a2 activation prevented oAβ-mediated synaptic depression in the hippocampus. Moreover, overexpression of Nr4a2 in the CA1 area of the hippocampus of APP_Sw,Ind_ mice ameliorated hippocampal-dependent spatial learning. These data support that a decrease in Nr4a2 function in the hippocampus is a factor involved in synaptic and learning impairment observed in the early stages of AD.

## MATERIAL AND METHODS

### Animals

C57BI-6J/RccHsd and APP_Sw,Ind_ (line J9; C857BI-6 background) mice were handled and used under institutional and national regulation approved by the Animal and Human Ethical Committee of the Universitat Autònoma de Barcelona (CEEAH 2896, DMAH 8787) following European Union regulation (2010/63/EU) and by the Albert Einstein College of Medicine Institutional Animal Care and Use Committee in accordance with the National Institute of Health guidelines. All animals were group-housed under standard laboratory conditions (food and water ad libitum, 22 ± 2°C, 12h light:dark cycle). We have used the C57BI-6J/RccHsd mice for primary neuronal cultures and to obtain acute hippocampal slices for electrophysiological studies. APP_Sw,Ind_ transgenic mice were used in behavioural studies. Both male and female animals were used in all the experiments.

### Human brain tissue

Postmortem human brain tissue analyzed in this study was provided by brain banks of Fundación CIEN (Instituto de Salud Carlos III, Madrid), Fundación Hospital Alcorcón (Madrid), Hospital Clínic-IDIBAPS (Barcelona) and Hospital Bellvitge (Barcelona). We analyzed hippocampal tissue samples from control subjects and patients diagnosed as Braak I-II (corresponding to presymptomatic AD stage), Braak III-IV (corresponding to mild cognitive impairment –MCI– due to AD) or Braak V-VI (corresponding to dementia due to AD). Information of sex, age and postmortem delay (PMD) is shown in supplemental table 1.

**Table 1.** Demographic and clinical information of human hippocampal tissue samples.

|  | **Control** | **Braak II** | **Braak III-IV** | **Braak V-VI** |
| --- | --- | --- | --- | --- |
| **Cohort size** | 13 | 13 | 13 | 28 |
| **Sex (W/M)** | 6/7 | 5/8 | 5/8 | 18/10 |
| **Age (years)** | 58.77 ± 12.87 | 70 ± 10.43 | 81.15 ± 12.77 | 81.5 ± 6.12 |
| Mean ± SD (range) | (43-79) | (52-86) | (46-98) | (67-92) |
| **PMD (h)** | 7.7 ± 4.23 | 5.46 ± 2.2 | 6.75 ± 3.24 | 8.4 ± 4.87 |
| Mean ± SD (range) | (2-16) | (3-9.75) | (2-14) | (2-20.5) |

Results are expressed as mean ± SD for each group, range (min-max). PMD, postmortem delay.

### Primary neuronal cultures

Primary hippocampal neurons were obtained from C57BI-6J/RccHsd mice pups (postnatal days 0-2). Neurons were mechanically and enzymatically dissociated with papain (Sigma) and plated (66,000 neurons/cm^2^) in Neurobasal-A medium supplemented with 2% B27 (Fisher Scientific) as previously described (Català-Solsona et al., 2023). Experiments were performed when hippocampal neurons were mature (18-21 DIV).

### Amyloid-β oligomerization

Lyophilised synthetic Aβ_1-42_ protein (Bachem or GenicBio) was kept at room temperature for 30 minutes and dissolved in cold hexafluor-2-propanol (HFIP) to obtain a concentration of 1 mM Aβ_1-42_. After homogenisation (1 to 3 h at room temperature), it was aliquoted in protein low-binding tubes and evaporated in a speed-vac at 800 x g <25°C for 20 min. The precipitate was stored at -80°C until use (< 6 months). The precipitated Aβ_1-42_ protein was dissolved in 100% dimethyl sulfoxide (DMSO) to get a stock of 5 mM. To allow oligomerization, Aβ_1-42_ was diluted to a final concentration of 100 μM by adding DMEM F-12 phenol-red free media (Life technologies) or artificial CSF (ACSF): 125 mM NaCl, 2.5 mM KCl, 33 mM D-glucose, 2 mM CaCl2, 1 mM MgCl2 and 25 mM HEPES (pH 7.3) for electrophysiology studies and kept 12 h at 4°C. Aβ_1-42_ oligomeric preparations were then biochemically analysed by BisTris/bicine western blotting using mouse monoclonal anti-APP 20.1 (provided by Dr. W.E. Nostrand) as primary antibody.

### Chemically induced LTP

Chemical LTP was induced as described previously (Otmakhov et al., 2004). After a 30 min incubation with ACSF at 37°C in a humidifier incubator with 95% O_2_ and 5% CO_2_, hippocampal cultures were treated with 50 μM forskolin, and 0.1 µM rolipram in ACSF with no MgCl_2_ during 10 min. Afterwards cultures were maintained for 3 hr and 50 min in complete ACSF before lysis. Amodiaquine (AQ) was added to the cultures 4 hr before cLTP when used.

### Immunoblotting

Primary neuronal cultures and tissue were homogenised in RIPA-lysis buffer containing (in mM): 50 Tris-HCl, pH 7.4, 150 NaCl, 2 mM EDTA, 0.5% Triton X-100, 1% NP-40, 0.1% SDS, 1 mM Na3VO4, 50mM NaF and 1 mM phenylmethylsulfonyl fluoride (PMSF) supplemented with cocktail of protease and phosphatase inhibitors (Sigma). Primary neuronal cultures were washed in PBS 1X and then lysed in cold RIPA lysis buffer by sonication using 30% of power (relative output 0.5) for 3 seconds (Sonic Dismembrator model 300, Dynatech) and samples were kept at -20°C until use. Human tissue was homogenated using a pestle homogeniser and solubilised 1h at 4°C. Then, lysates were sonicated three times for 15 seconds and centrifuged at 9,300 x g for 15 minutes at 4°C. The supernatant was kept at -80°C until use. Proteins were separated on 7.5-12% SDS-PAGE and transferred to nitrocellulose membranes (GE Healthcare). Blots were blocked 1 h with 10% dry milk, 0.1% BSA pH 7.4 in phosphate buffered saline (PBS) or Tris-HCl buffered saline (TBS) and incubated at 4°C overnight with primary antibodies (anti-Nr4a2, Abcam, ab41917, 1:500 dilution; anti-GluA1, Merck-Millipore, AB1502, 1:1000 dilution; anti-phosphoSer845GluA1, Abcam, ab76321, 1:1000 dilution; anti-GAPDH, Thermo Fisher Scientific, AM4300, 1:10000 dilution). After incubation with appropriate horseradish peroxidase-conjugated secondary antibodies (HRP-linked anti-mouse or anti-rabbit IgG; BD Biosciences) at RT for 1 h, blots were developed using ECL^TM^ Western blotting Detection Reagents (GE Healthcare). Blots densitometry was performed using ImageJ (National Institutes of Health, Bethesda, MD), and protein levels were corrected for corresponding loading control.

### Lentiviral vectors production

A construct to overexpress Nr4a2 was generated with the In-Fusion HD Cloning Kit (Clontech) following supplier’s recommendations. FUGWshNr4a2 vector was digested using AgeI restriction enzyme (New England BioLabs) and pWpi vector using PmeI restriction enzyme (New England BioLabs). 50 μl of the digested vector was resolved in 1% agarose-TAE gel and DNA band was cut and purified using the NucleoSpin Gel and PCR clean-up kit as previously. Nr4a2 coding sequence was obtained from pcDNA3.1Nr4a2V5HisB plasmid kindly provided by Dr. Ángel Juan García Yagüe (Instituto de Investigaciones Biomédicas “Alberto Sols”, Madrid, Spain). Specific primers following In-Fusion ND Cloning kit recommendations were used to amplify Nr4a2 coding sequence. FUGWshNr4a2+Nr4a2 forward: 5’-ATCCCCGGGTACCGGATGCCTTGTGTTCAGGCGCAGTATG-3’; FUGWshNr4a2+Nr4a2 reverse: 5’-CATGGTGGCGACCGGCGTAGAATCGAGACCGAGGAGAG-3’. The In-Fusion reaction mixture were transformed into Stellar Competent Cells (Clontech) using the heat shock Method.

Human embryonic kidney 293T (HEK293T) cells were grown in supplemented DMEM medium (Thermo Fisher Scientific). At 70% of confluence, HEK293T cells were transfected using the CalPhos Mammalian Transfection kit (Clontech) following supplier’s recommendations. In each dish of 100 mm of diameter, the DNA mixture consisted in 20 μg of specific DNA and 10 μg each of psPax2 and pMD2.G plasmids (from Dr. Didier Trono, Addgene) and 250 mM CaCl2 (in a final volume of 500 ul) mixed drop wise with 2X HBS. After 20 minutes at room temperature, the DNA mixture solution was added drop wise to HEK293T cells and incubated for 8 hours at 37°C in a humidifier incubator with 5%CO2 / 95%air. Then, medium was replaced by fresh supplemented DMEM. HEK393T culture medium was collected 24, 36 and 48 hours after transfection and filtered through a 0.45 μm pore size filter to remove cellular debris. Then, medium was spun at 47,000 x g for 2 hours at 4°C to concentrate the lentiviral vector particles, which were resuspended in 100 μl cold PBS 1X without supplements and kept at 4°C overnight with soft shaking before being aliquot and stored at -80°C. Biological titers of the viral preparations were assessed by transducing HEK293T cells with serial dilutions and checking the percentage of green fluorescent protein (GFP)-positive cells by flow cytometry (Cytomics FC 500, Beckman Coulter). Hippocampal-cultured neurons were transduced at 7DIV with lentiviral vectors (1-2 transducing units/cell).

### Hippocampal slice preparation

Acute transverse hippocampal slices were prepared from P25 to P40 C57BL/6J mice of either sex (400 µM) using a VT1200 Leica vibrate. The cutting solution contained the following (in mM): 93 N-Methyl-d-glucamine, 2.5 KCl, 1.25 NaH2PO4, 30 NaHCO3, 20 HEPES, 25 D-glucose, 2 Thiourea, 5 Na-Ascorbate, 3 Na-Pyruvate, 0.5 CaCl2, 10 MgCl2. The ACSF recording solution contained (in mM): 124 NaCl, 2.5 KCl, 26 NaHCO3, 1 NaH2PO4, 2.5 CaCl2, 1.3 MgSO4 and 10 D-glucose. Slices were transferred directly to ACSF at 34°C in warm water bath and then incubated at RT for at least 1 h. All solutions were equilibrated with 95% O2 and 5% CO2 (pH 7.4).

### Electrophysiology

Extracellular field potentials (fEPSPs) were recorded with a patch-type pipette filled with 1M NaCl and placed in the stratum radiatum (50-100 µm from CA1 pyramidal neurons somas). Experiments were performed at 29 ± 1°C in a submersion-type recording chamber perfused at 2 ml/min ACSF containing 100 µM picrotoxin and, when indicated, oAβ (0.5 µM) and/or amodiaquine (AQ; 10 µM). Low-frequency stimulation (LFS; 300 pulses at 1 Hz) was induced after 1 hr of stable baseline and fEPSPs were monitored at least for 45 min.

Long-term potentiation (LTP) was induced after 45-60 min of stable baseline by theta burst stimulation (TBS), which consisted in a series of 10 bursts of 5 stimuli (100 Hz within the burst, 200 ms interburst interval) repeated 4 times (5 s apart). When AQ was used, slices were incubated in the dark with AQ for 2 hr prior to transfer to the recording chamber and fEPSPs recordings were also performed in the dark. The magnitude of LFS or LTP was determined by comparing baseline-average responses before induction with the last 15 min of the experiment.

### AAV production and stereotaxic surgery

Adeno-associated viral vectors (AAV) were generated by the Vector Production Unit (Centre de Biotecnologia Animal i de Teràpia Gènica – CBATEG –, Universitat Autònoma de Barcelona). AAV2/10 to overexpress Nr4a2 (containing Nr4a2.V5) under the CMV promoter, were generated. For stereotaxic surgery, C57BL/6J 4.5-month-old mice were first anaesthetised in a chamber with 3-5% isoflurane. They were placed in a stereotaxic frame (Kopf) with the head subjected to blunt ear bars and the nose placed into a ventilator and anaesthesia system using continuous isoflurane (up to 5% for induction and 1.5-3% for maintenance in 4.5-month-old mice).

Either AAV2/10.H1.scramble.RSV.GFP or AAV2/10.H1.shNr4a2.RSV.GFP were injected (1 uL; 6x10^12^ gc/ml; 0.15 μl/min). Injections were performed bilaterally into the dorsal CA1 region (2.18 mm posterior to bregma, 1.75 lateral to bregma, 1.6 ventral from dural surface, according to Paxinos and Franklin mouse brain atlas) using a beveled needle (Hamilton). Both male and female mice were used with similar ratio for the two types of viruses. Slices for electrophysiology were prepared from NMDG-perfused injected animals 3-4 weeks after injection.

### Behaviour

All mice were housed under standard laboratory conditions at the animal facility of the Universitat Autònoma de Barcelona, on a 12 hour light/dark cycle, 22 ±2°C, relative humidity 50-60% and with food and water available ad libitum. Littermates were housed together (maximum 5 mice per cage) until one week before starting the behavioural tests, when they were separated into individual cages to avoid fighting. A total number of 80 WT (non-transgenic) and APP_Sw,Ind_ mice were used (40 each genotype; 50/50 male and female). Viral stereotaxic hippocampal injections to overexpress Nr4a2 were performed at 4.5-month-old mice, and behaviour and cognition were evaluated at 6 months of age.

A circular pool for mice (PanLab; 120 cm diameter, 84 cm height, 23°C opaque water) with a hidden platform (11 cm diameter, 1 cm below the water surface) was used for the Morris water maze test. Mice were gently released facing the wall of the pool from one randomly selected point (one different each day) and allowed to swim for 1 minute to find the platform. The first day (cue learning task), the platform was in the middle of the N-E quadrant. It was visible 1 cm above the water surface and cued with a visible flag. In the next days (place learning task acquisition phase), the platform was hidden and located in a reversed position (in the middle of the S-W quadrant). We placed four geometric figures on each wall of the room that were used as external visual clues. In all trials, mice reaching the platform were left there for 10 seconds, and mice failing to find the platform were placed on it for the same time. For all learning days, variables regarding the time (escape latency), distance and speed were recorded by a computerised tracking system (ANY-maze).

In the Novel Object Recognition test (NOR), mice were placed individually in the center of an open field for 1 minute (habituation to context). Afterward, mice were exposed to two identical objects (sample) and allowed to explore them (defined as nose facing the oat 2 cm, sniffing, or biting the objects) until reaching criteria (a total accumulated time of 20 seconds) during a period of a maximum of 10 minutes (training). After 4 hours, mice were subjected to a retention test in a 5 minute session with two objects: one identical to those previously explored (familiar) and a new one (novel). The time that the mice spent exploring each of the objects was measured and preference for the novel object was expressed as a discrimination index (DI): DI = (time novel – time familiar) / (time novel + time familiar) x 100%. Mice that explored both objects for less than three seconds during training or two seconds during testing were not considered for the analysis.

### Statistical analysis

Statistical analysis was performed using GraphPad Prism software v6.01 (GraphPad Software Inc., California, USA) and SPSS 17.0 software for behavioral studies.

We performed either unpaired Student’s *t*-tests or analyses of variance (ANOVAs; one-, two-or three-way) followed by appropriate between-group comparisons according to each analysis requirements, Bonferroni or Tukey *post hoc* test for parametric samples or two-sided nonparametric Mann-Whitney test for nonparametric samples.

For behavioral analysis, differences between genotype and treatment interactions were analyzed by repeated-measures ANOVA, and correlation analysis was performed between different variables using nonparametric Spearman correlation coefficient.

Data is shown as the mean ± standard error of the mean (SEM) or standard deviation (SD). Statistically significant difference was set at *p*-value < 0.05 and is indicated as follows: *p < 0.05, **p < 0.01, ***p < 0.001, ****p < 0.0001.

## RESULTS

### Activity-dependent induction of Nr4a2 is impaired by soluble forms of Aβ peptide

It is now widely accepted that oAβ are one of the initial mediators leading to early synaptic dysfunction in AD (Forner et al., 2017). Since we have previously reported that Nr4a2 has a key role in hippocampal synaptic plasticity (Català-Solsona et al., 2023), we wanted to address the eventual involvement of Nr4a2 in the synaptic failure occurring at early AD stages. For that purpose, we firs addressed whether oAβ could alter Nr4a2 levels in hippocampal neurons (Fig. 1). We have observed a significant reduction in the activity-dependent increase of Nr4a2 protein levels in mature hippocampal cultures in the presence of oAβ (45% and 35% reduction after 2 and 4 hours of oAβ incubation, respectively; Fig. 1B). The decrease observed in Nr4a2 protein levels in the presence of oAβ was not found in Nr4a2 mRNA since the bicuculline-mediated increase in Nr4a2 mRNA was not affected by oAβ (Fig. 4C).

**Figure 1.**
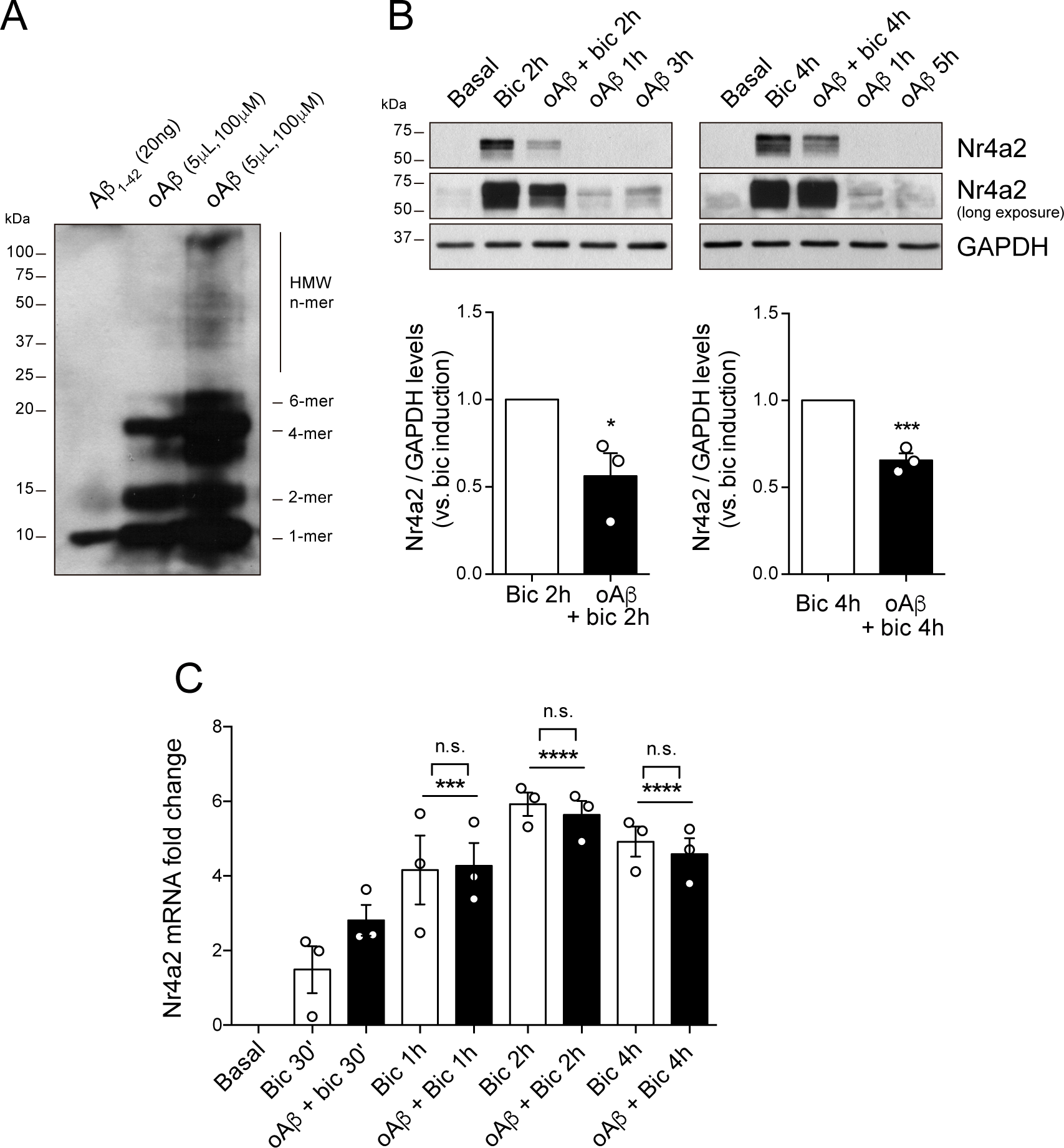
oAβ reduce the activity-dependent increase in Nr4a2 protein levels in mature hippocampal neurons. (A) Representative western blotting image of Aβ species present in soluble monomeric Aβ1-42 and oligomeric preparations. (B) Nr4a2 protein and (C) Nr4a2 mRNA levels after bicuculline (bic; 50μM) treatment in the presence or absence of oAβ (5μM) applied 30 minutes before bic treatment. Data represent mean ± S.E.M. n=3 independent hippocampal cultures. Statistical analysis in B was determined by Student’s unpaired two-tailed t-test and in C by one-way ANOVA followed by Tukey *post hoc* test. *p<0.05, ***p<0.001, ****p<0.0001.

### Nr4a2 protein levels are decreased in human postmortem hippocampal tissue at early AD stages

Nr4a2 expression has been previously examined in AD patients with contradictory results (Parra-Damas et al., 2014; Annese et al., 2018; Moon et al., 2019). Here, we have checked Nr4a2 protein levels in postmortem hippocampal tissue samples from control subjects and AD patients, and we found a significant decrease in Nr4a2 protein levels in Braak II stage (30% decrease), corresponding to a pre-symptomatic stage where synaptic deficits are already occurring, compared with healthy age-matched controls (Fig. 2). By contrast, no changes were observed at Braak III-IV and V-VI stages when compared with controls.

**Figure 2.**
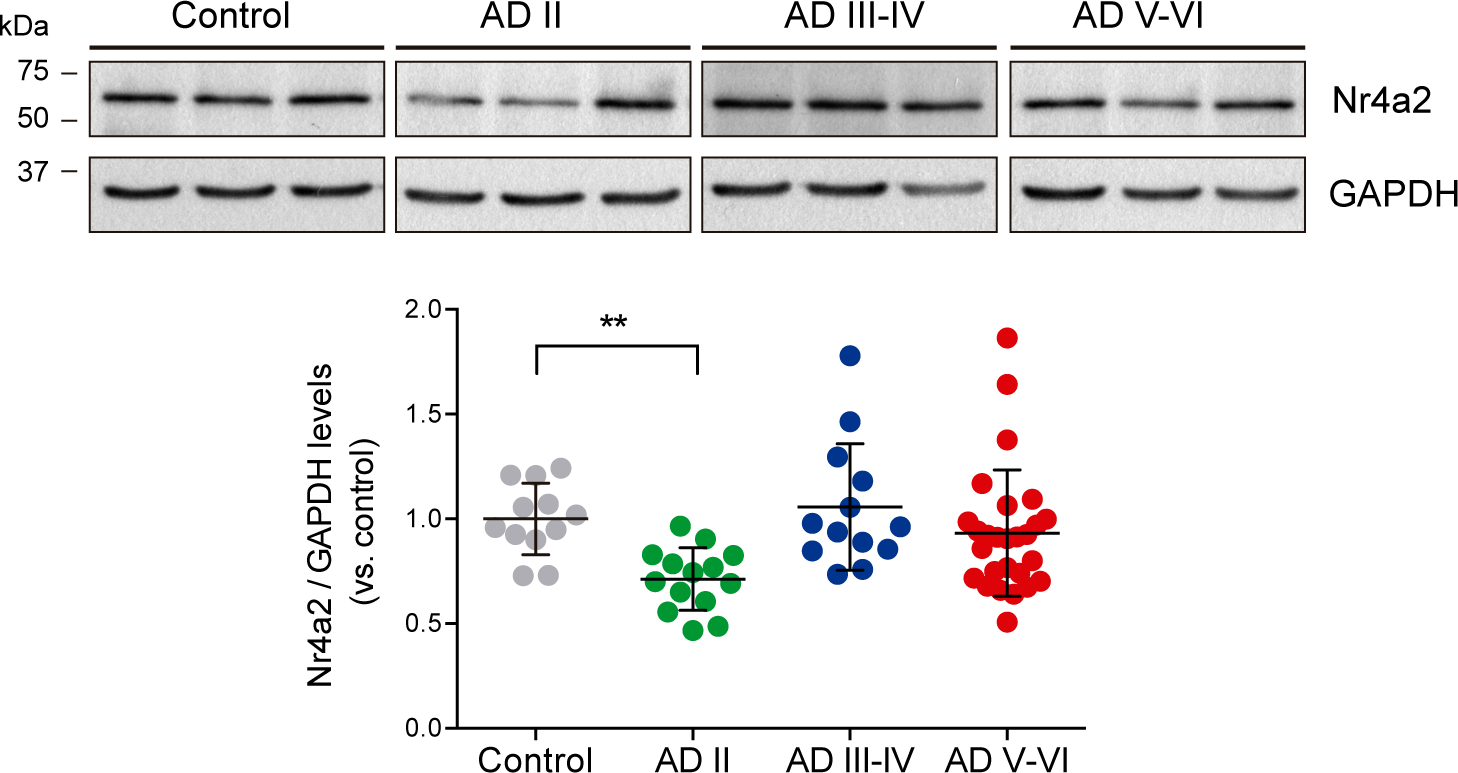
Nr4a2 levels in AD postmortem human hippocampus. Representative western blotting images and quantification of Nr4a2 protein levels in hippocampal human postmortem tissue samples of control subjects and AD patients classified in different Braak stages. n=13-28/group. Data represent mean ± S.E.M. Statistical analysis was determined by Kruskal-Wallis test. **p<0.01.

### Nr4a2-activation rescues the oAβ-mediated synaptic depression observed in mature hippocampal neurons

oAβ disrupts hippocampal synaptic plasticity impairing NMDA-dependent LTP and promoting LTD (Walsh et al., 2002; Hsieh et al., 2006; Shankar et al., 2008; Li et al., 2009). A mechanism involved in this effect is the oAβ-mediated decrease of surface expression of AMPARs (Hsieh et al., 2006; Minano-Molina et al., 2011). Since, we have recently reported that Nr4a2 activation increases postsynaptic AMPARs and effectively blocks LTD (Català-Solsona et al., 2023), we addressed whether Nr4a2 activation was able to rescue the oAβ-mediated deficits in hippocampal synaptic plasticity. We used the Nr4a2 activator amodiaquine (AQ) to test the effect of Nr4a2 activation on cLTD in primary cultures of hippocampal neurons (Fig. 3A). AQ increased Ser845-GluA1 phosphorylation in basal conditions and completely rescued the oAβ-mediated dephosphorylation at Ser845-GluA1 (phosphorylated levels of Ser845-GluA1-AMPARs after AQ treatment: 1.9 ±0.21, oAβ + AQ: 1.66 ±0.3, oAβ: 0.6 ±0.07 fold change vs. basal). The rescue effect of AQ vs oAβ-mediated synaptic depression was also observed in hippocampal slices (Fig. 3B). Whereas subthreshold low-frequency stimulation (LFS, 300 pulses, 1 Hz) didn’t trigger a significant LTD, the presence of oAβ (500 nM) induced a robust LTD facilitation. Bath application of AQ (3 hrs - 10 µM) blocked oAβ-mediated synaptic depression. oAβ has also been described to impair LTP in the hippocampus (Walsh et al., 2002; Shankar et al., 2008). Therefore, we proceed to test whether Nr4a2 activation could also rescue from oAβ-mediated blockade of hippocampal LTP. We applied a cLTP protocol consisting in treating mature cultures of hippocampal neurons with forskolin (50 µM) and rolipram (0.1 µM) for 10 min in Mg^2+^-free ACSF (Fig. 4A-B) that resulted in prolonged NMDAR-dependent LTP and increases phosphorylation of GluA1 at Ser-845 (Zwain and Yen, 1999; Otmakhov et al., 2004; Oh et al., 2006). As could be seen in Fig. 3A, the cLTP-mediated increase in pSer^845^GluA1 was significantly reduced in the presence of oAβ and could be rescued by incubation with the Nr4a2 activator AQ (cLTP: 7.1 ± 0.46; cLTP+ oAβ: 4.4 ± 0.5; cLTP+ oAβ+AQ: 7.8± 0.5-fold change vs basal). Importantly, AQ was not only able to restore the oAβ-mediated impairment of cLTP but also increased, around 30%, the Ser^845^GluA1-AMPAR phosphorylation after a cLTP protocol (Fig. 3A). Similar results were obtained in mature hippocampal cultures that were transduced at 7 DIV with lentiviral vectors to overexpress Nr4a2 (Fig 3B). Overexpression of Nr4a2 efficiently blocked the oAβ-mediated decrease in pSer^845^GluA1 after the cLTP protocol. Furthermore, we investigated whether AQ blocks oAβ-mediated effect on LTP in acute hippocampal slices after a theta burst stimulation (TBS) of Schaeffer collaterals. A reduction on LTP was observed in oAβ-treated slices (Fig 4C). When AQ was associated to oAβ, TBS induced a sustained increase in the fEPSP slope when compared to oAβ-treated slices. Altogether our data indicate that Nr4a2 activation can rescue oAβ-mediated impairment of hippocampal synaptic plasticity.

**Figure 3.**
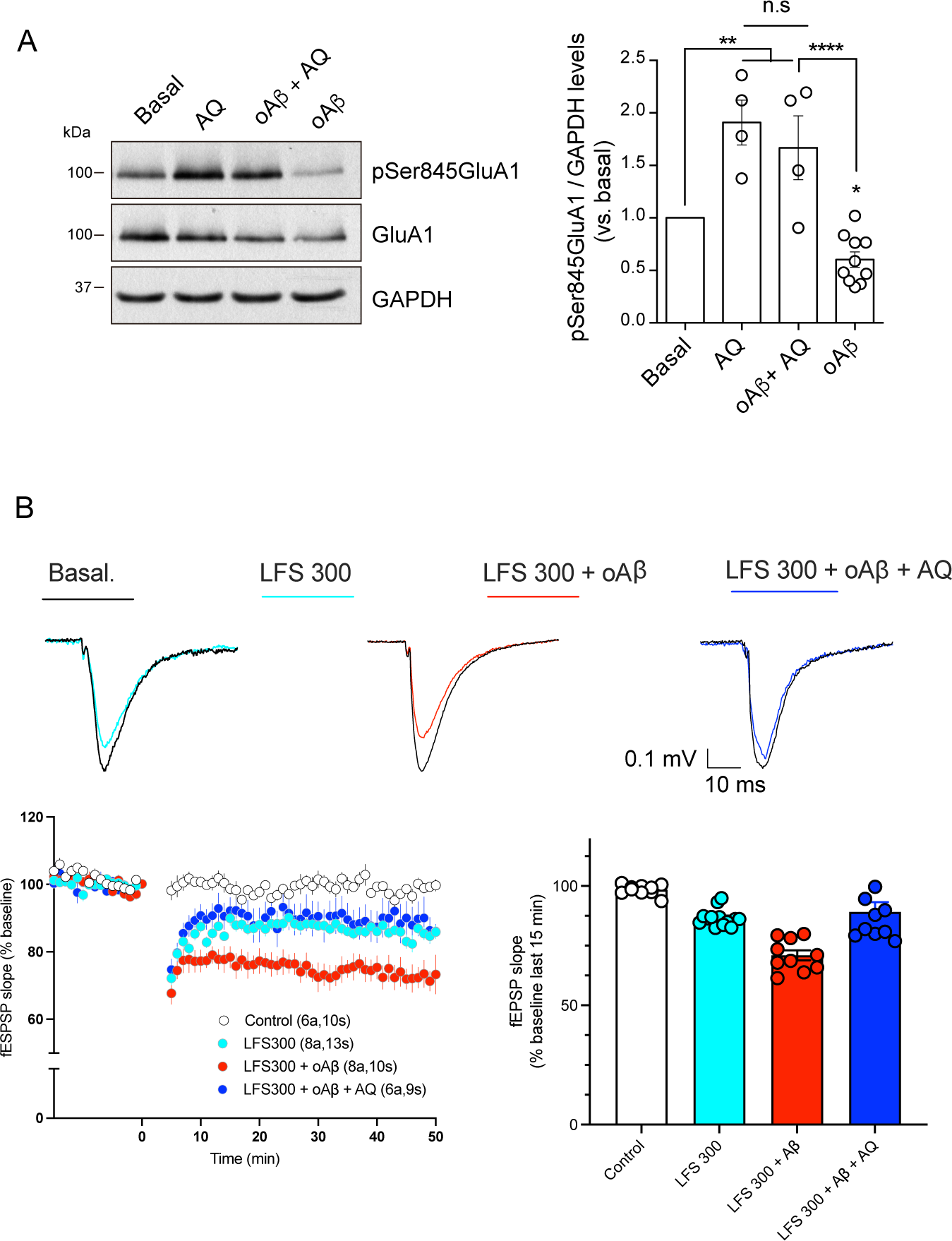
Nr4a2 rescues the oAβ-dependent synaptic depression in the hippocampus. (A) Total and Ser845-phosphorylated GluA1-AMPAR protein levels in oAβ-treated hippocampal neurons in the presence or absence of amodiaquine (AQ; 10μM). n=4 independent hippocampal cultures. (B) fEPSP responses of Schaffer collateral inputs in acute hippocampal slices, treated with oAβ (0.5 µM) and AQ (10 µM), before and after LFS (300 pulses at 1 Hz). Traces represent averages of 25 responses. Number in parentheses indicate the number of animals (a) and slices (s). Data represent mean ± S.E.M. Statistical analysis was determined one-way ANOVA followed by Tukey *post hoc* test. *p<0.05, **p<0.01, ****p<0.0001.

**Figure 4.**
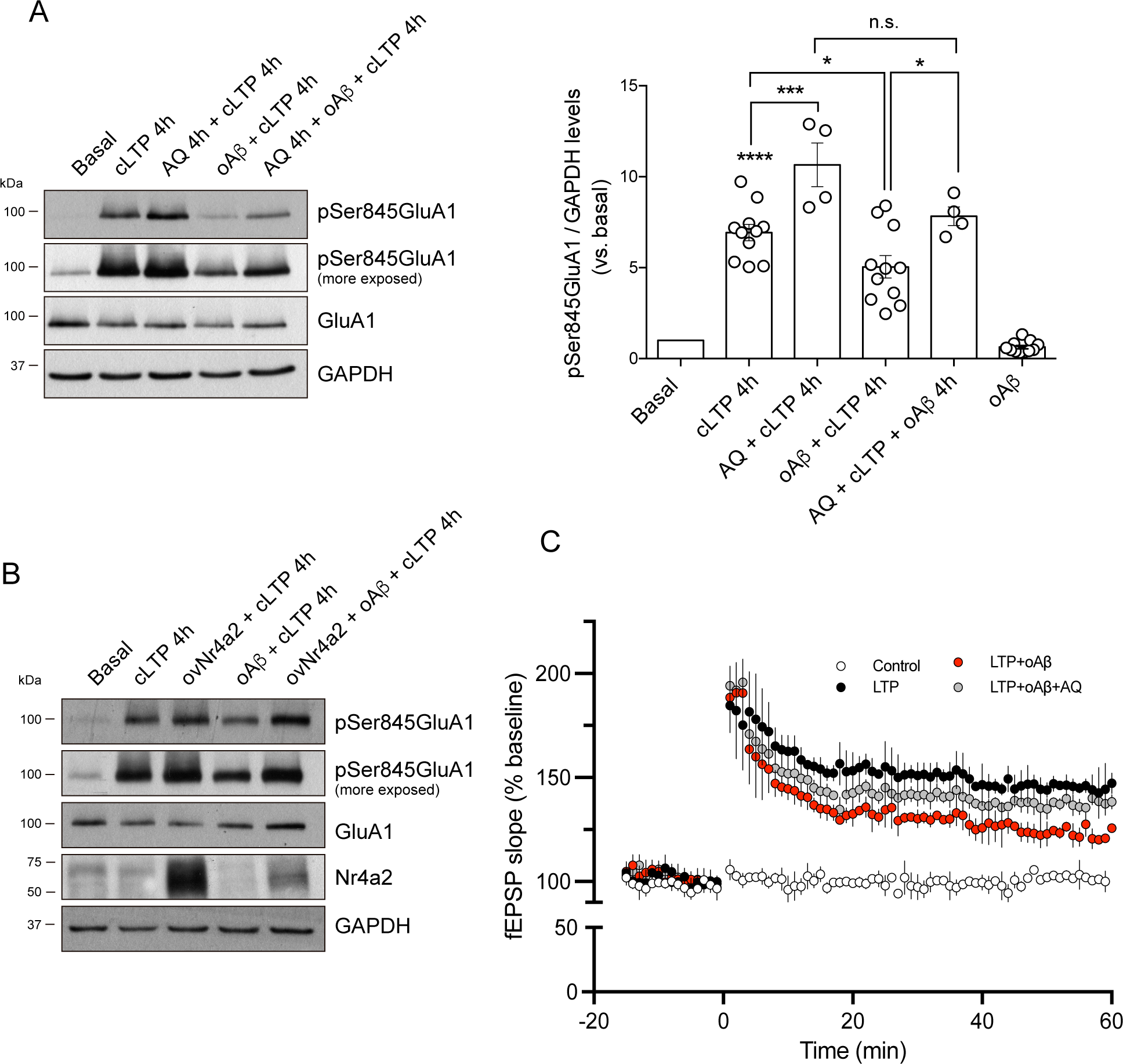
Nr4a2 potentiates cLTP and rescue oAβ-mediated cLTP impairment in mature hippocampal neurons. (A) Ser845-phosphorylated GluA1-AMPAR protein levels after cLTP protocol in the presence or absence of the Nr4a2 agonist amodiaquine (AQ; 10 µM) and/or oAβ (5 µM). n=4 independent hippocampal cultures. (B) Total and Ser845-phosphorylated GluA1-AMPAR protein levels after cLTP protocol in hippocampal cultures transducer with a lentiviral vector to overexpress Nr4a2. n=3 independent hippocampal cultures. (C) fEPSP responses of Schaffer collateral inputs in acute hippocampal slices, treated with oAβ (0.5 µM) and AQ (10 µM), before and after LTP induction. Traces represent averages of 25 responses. Numbers in parentheses indicate the number of animals (a) and slices (s). Data represent mean ± S.E.M. Statistical analysis was determined by ANOVA followed by Tukey *post hoc* test. *p<0.05, **p<0.01, ****p<0.0001.

### Nr4a2 hippocampal overexpression ameliorates the cognitive deficits observed in the APP_Sw,Ind_ mouse model of AD

Our results show that Nr4a2 activation blocks hippocampal LTD (Català-Solsona et al., 2023) and oAβ-mediated synaptic depression and impairment of LTP (Fig. 3 and 4). LTP and LTD are synaptic processes related to types of learning and memory (Lynch, 2004; Neves et al., 2008; Collingridge et al., 2010; Nicoll, 2017). Thus, we wanted to study whether Nr4a2 hippocampal overexpression could ameliorate the early learning deficits observed in the APP_Sw,Ind_ mouse model of AD (España et al., 2010). We overexpressed full-length Nr4a2 construct by stereotaxic AAV injections in the CA1 of 4.5 month-old WT and APP_Sw,Ind_ mice. We used AAV2/10 serotype, which has been characterised by high and specific gene transduction in neurons of adult mice brain (Cearley and Wolfe, 2006). Behaviour was assessed at 6 months of age. To analyse spatial learning, a key component of episodic memories that is hippocampus-dependent and that is affected in AD, mice were subjected to the MWM test as described in Material and Methods section.

The distance covered to locate the platform in the cue learning task and the first two days of the acquisition phase are shown in Figure 5. Learning acquisition was slower in APP_Sw,Ind_ mice, covering more distance to reach the platform compared to WT mice, and was significantly restored by Nr4a2 overexpression, mimicking the same distance covered by the WT group to reach the platform the second day of the acquisition phase (Fig. 5A; distance in cm to reach the platform; WT: 489.4 ±56.6, WT-Nr4a2: 431.7 ±65.7, APP_Sw,Ind_: 661.9 ±93, APP_Sw,Ind_-Nr4a2: 431.89 ±37). In the probe trial, APP_Sw,Ind_ mice crossed less times the place where the platform was located, and APP_Sw,Ind_-Nr4a2 mice crossed the same number of times as the WT group (Fig. 5C), showing an effect of Nr4a2 overexpression not only in the learning, but also in the spatial memory. Furthermore, to check that hippocampal overexpression of Nr4a2 was not affecting non-hippocampal dependent memories, mice were subjected to the NOR test that does not require dorsal hippocampus for retrieval (Vogel-Ciernia et al., 2013). No significant differences in total exploration time (Fig. 6B) and preference for the two objects (Fig. 6C) were observed at training.Conversely, all four groups showed preference for the new object in the probe test with no differences between genotype or Nr4a2 overexpression showing no ORM deficits in this cohort of APP_Sw,Ind_ mice and no effect of Nr4a2 hippocampal overexpression (Fig. 6D).

**Figure 5.**
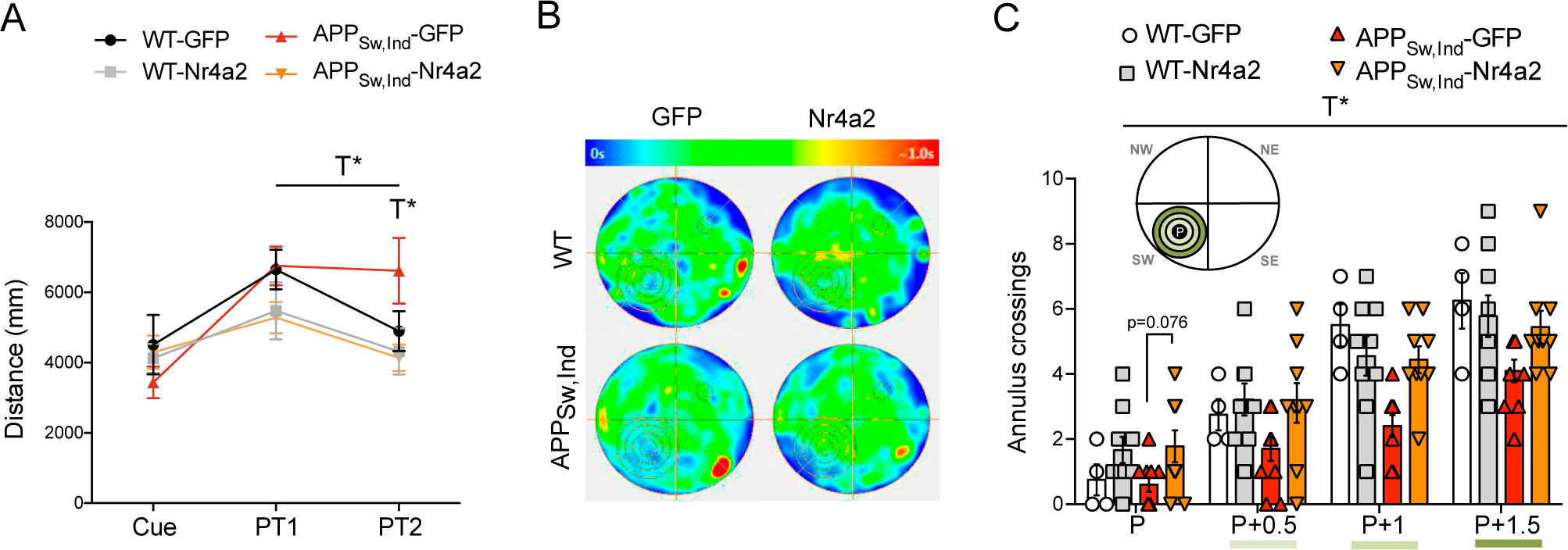
Nr4a2 hippocampal overexpression ameliorates spatial learning deficits in APP_Sw,Ind_ mice. AAV-mediated Nr4a2 hippocampal overexpression was performed as described in Methods. (A) Distance travelled in the Morris Water Maze (MWM) test to reach the platform in the cue and place learning task 1 and 2 of 6 month-old WT and APP_Sw,Ind_ mice with or without Nr4a2 overexpression. (B) Cumulative swim track representations (heat maps) of each group of mice during the probe trial of the MWM test. Relative occupancy values are indicated by the colour code (see colour bar; red denotes high occupancy, blue denotes low occupancy values). (C) Annulus crossings through the platform in the probe trial of 6 month-old WT and APP_Sw,Ind_ mice with or without Nr4a2 overexpression. n= 8-10 mice/group. Data mean ± S.E.M. Statistical analysis was determined by three-way ANOVA followed by Tukey *post hoc* test. *p<0.05. T, treatment effect

**Figure 6.**
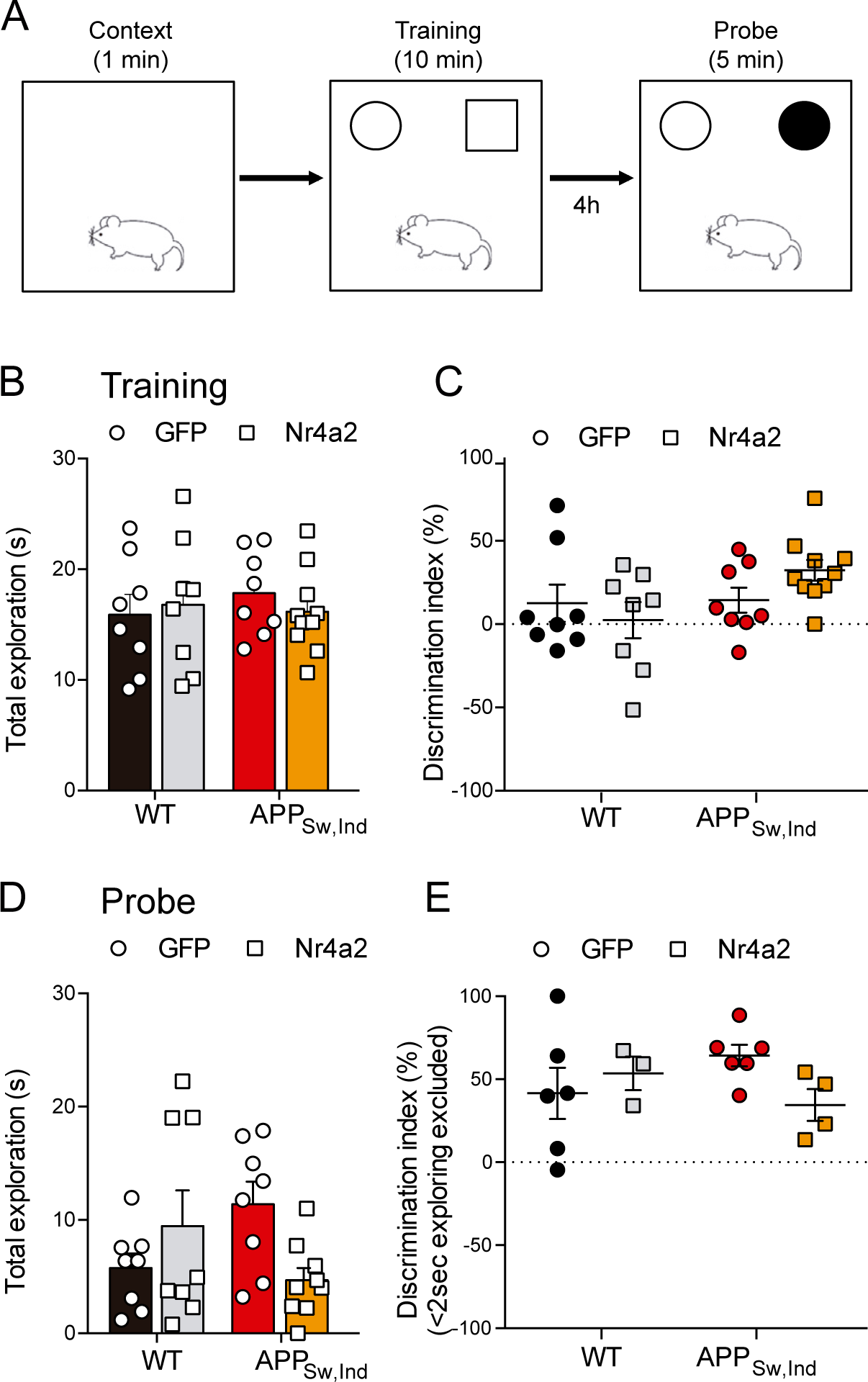
Nr4a2 hippocampal overexpression does not affect object recognition memory. (A) Design of the novel object recognition (NOR) test used in the study. (B-E) Total exploration and discrimination index in the training (B,C) and probe (D,E) of the NOR test in 6-month-old WT and APP_Sw,Ind_ mice with or without Nr4a2 overexpression. n= 8-10 mice/group. Data mean ± S.E.M. Statistical analysis was determined by two-way ANOVA followed by Tukey *post hoc* test.

## DISCUSSION

The hippocampus, a brain area critical for learning and memory, is especially vulnerable to damage at early stages of AD (Scheff et al., 2007; La Joie et al., 2013) and seems to be the most severely affected brain area by the loss of synaptic proteins (Claudia et al., 2011). Indeed, hippocampal synaptic loss is currently established as the best neurobiological correlate of the cognitive decline observed in human AD patients and reflects the synaptic dysfunction that initiates the disease process (Terry et al., 1991; DeKosky et al., 1996; Bereczki et al., 2016). AD early cognitive impairments have been attributed to an oAβ-induced disruption of synaptic plasticity. Nevertheless, the underlying molecular mechanisms are yet to be fully understood. A substantial body of evidence indicates that activity-regulated transcriptional networks promote the refinement and plasticity of neuronal circuits, which are essential for cognitive functions (Alberini, 2009; Benito and Barco, 2015). Among them, several reports have described that some members of the Nr4a family of orphan receptors, especially Nr4a2, are involved in hippocampal synaptic plasticity and memory processes (Hawk et al., 2012; Bridi and Abel, 2013; Català-Solsona et al., 2021) by a molecular pathway that regulates synaptic AMPARs (Català-Solsona et al., 2023). Down-regulation of activity-dependent genes involved in synaptic plasticity and memory are associated with early learning and memory deficits in mouse models of AD (Saura and Valero, 2011; Parra-Damas et al., 2014; Marcello et al., 2018). A decrease in Nr4a2 expression is observed in APP_Sw,Ind_ mice when initial hippocampal-dependent spatial memory deficits coincide with initial oAβ accumulation (España et al., 2010). Since Nr4a2 is involved in AMPARs modulation (Català-Solsona et al., 2023) and oAβ has been shown to promote synaptic depression by removal of synaptic AMPARs (Hsieh et al., 2006; Minano-Molina et al., 2011), we wanted to study the eventual role of Nr4a2 in AD-mediated synaptic and cognitive dysfunction. Our results show that oAβ blocked the activity-mediated increase in Nr4a2 protein levels in mature hippocampal neurons. By contrast, the activity-mediated increase of Nr4a2 mRNA levels was not altered. Our results with oAβ differ from the ones reported in cerebellar granule cells with Aβ_1-42_ fibrils (Terzioglu-Usak et al., 2017). Besides the distinct cell type used in both studies, these observations open the possibility that diverse Aβ structures could have different effects on Nr4a2 and other proteins levels. In any case, our results suggest that oAβ could affect activity-mediated increase in Nr4a2 protein levels post-transcriptionally, probably by promoting Nr4a2 degradation. In fact, we have previously described (Català-Solsona et al., 2023) that Nr4a2 levels in basal conditions are highly regulated by proteasome activity.

Nr4a2 expression has also been previously examined in AD human brains without achieving consistent results. Increased hippocampal Nr4a2 mRNA levels were previously reported in a cohort of late-onset AD patients (Annese et al., 2018). Conversely, decreased levels of Nr4a2 mRNA expression at Braak III-IV and V-VI stages compared to controls were found in postmortem human hippocampus (Parra-Damas et al., 2014). Likewise, reduced Nr4a2 protein levels were also reported in the hippocampus and the superior frontal cortex of AD patients (Moon et al., 2019). These discrepancies between studies could arise from the distinct demographic and clinical features of the samples, including different gender, age, origin, social status, or sample processing. We found a significant decrease of Nr4a2 protein levels specifically at Braak II stage, corresponding to preclinical AD, where symptoms are still absent albeit synaptic deficits have already started. It is also important to note that no differences were observed in Nr4a2 protein levels in Braak III-IV and Braak V-VI compared to controls because, although most samples from these stages had less Nr4a2 protein levels compared to healthy control subjects, few expressed high amounts of Nr4a2. Therefore, to conclude whether Nr4a2 protein levels are indeed decreased at late stages of AD, more samples should be analyzed.

Numerous studies have demonstrated that oAβ alter excitatory synaptic transmission and cause synapse loss (Hsia et al., 1999; Lennart et al., 2000; Shankar et al., 2007; Mucke and Selkoe, 2012). It has been reported that oAβ impair LTP (Hsieh et al., 2006; Shankar et al., 2008) and facilitates LTD (Walsh et al., 2002; Hsieh et al., 2006; Sanderson et al., 2021) in the hippocampus, likely due to promote receptor internalisation of inotropic glutamate receptors (Snyder et al., 2005; Hsieh et al., 2006; Minano-Molina et al., 2011) and excessive activation of extra synaptic NMDA receptors (Li et al., 2011). Moreover, it is believed that oAβ-mediated synaptic depression shares similar (if not equal) mechanisms to LTD elicited by low-frequency stimulation (LFS) of 900 pulses at 1 Hz. Since we have recently reported that pharmacological activation of Nr4a2 with amodiaquine produced a blockade of LFS-mediated LTD in CA1 and Nr4a2 knock-down in CA1 increased LFS-LTD (Català-Solsona et al., 2023), we tested in the present study whether pharmacological activation of Nr4a2 was able to revert the effect of oAβ on hippocampal synaptic plasticity. Our results show that pharmacological Nr4a2 activation with amodiaquine blocked both the oAβ-mediated cLTD-like dephosphorylation of GluA1Ser845 (Ehlers, 2000; Minano-Molina et al., 2011) in mature hippocampal cultures and oAβ-facilitation of synaptic depression in acute hippocampal slices. Also, Nr4a2 activation partially reversed: a) the oAβ-mediated blockade of the extrasynaptic delivery of AMPARs mediated by chemical LTP; b) the observed reduction of LTP in acute hippocampal slices. Thus, pharmacological activation of Nr4a2 is a feasible way to modify the impact of oAβ on hippocampal synaptic plasticity.

AD mouse models usually display learning and memory impairment that develops differently depending on the progression of the Aβ-like pathology in each specific mouse model. The APP_Sw,Ind_ mouse model phenotype is characterized by progressive Aβ pathology, showing increased Aβ_1-42_/ Aβ_1-40_ ratio as early as at 3 months of age (Mucke et al., 2000), coinciding with impaired hippocampal basal synaptic transmission and plasticity (Saganich et al., 2006). At 6 months-old, APP_Sw,Ind_ mice have CRE-transcriptional deficits (Espana et al., 2010; España et al., 2010), decreased pSer845-GluA1-AMPAR levels associated to early spatial memory deficits (Minano-Molina et al., 2011) and increased anxiety (España et al., 2010b). At this age, hippocampal gliosis is also present (Wright et al., 2013), with diffusive Aβ plaques in the DG and neocortex, which become more evident from the age of 9 months (Mucke et al., 2000). Using this model, we overexpressed Nr4a2 in the CA1 hippocampal subregion by using stereotaxic injections of AAV containing the full length Nr4a2 and learning and memory was assessed at 6 months of age mimicking early stage of the pathology. Previous studies have implicated Nr4a2 in long-term spatial information storage in the hippocampus. It is known that Nr4a2 mRNA is increased in CA1 and CA3 hippocampal subregions after a spatial food search task (Peña de Ortiz et al., 2000). Likewise, hippocampal infusions of Nr4a2 antisense oligodeoxynucleotides impair learning and long-term memory in a holeboard spatial discrimination task in rats (Colón-Cesario et al., 2006). In agreement, we found that Nr4a2 overexpression in hippocampal CA1 significantly ameliorated spatial learning and memory deficits of the APP_Sw,Ind_ mice in the Morris water maze test. Similarly, previous studies reported that pharmacological activation of Nr4a2 improved performance in the Y-maze of 5XFAD mice and in a spatial object recognition tasks in aged mice (Moon et al., 2019; Chatterjee et al., 2020). On the other hand, some discrepancies exist about the possible involvement of Nr4a2 in non-hippocampus-dependent memories. A recent study has demonstrated that hippocampal overexpression of Nr4a2 did not ameliorate object recognition memory deficits, a hippocampus-independent task (Kwapis et al., 2019). Nevertheless, intraperitoneal injections of a Nr4a2-specific agonist enhanced long-term memory in the NOR test (Kim et al., 2016), suggesting that this transcription factor could also be enhancing non-hippocampus-dependent memories. We have not found significant differences in total exploration time and preference for the objects during training in the NOR test. Furthermore, all groups tested showed preference for the new object in the probe test with no differences between genotype or treatment indicating that the cohorts of APP_Sw,Ind_ mice utilized in this study did not show deficits in object recognition memory, limiting our abilities to reach a conclusion on the potential effects of Nr4a2 hippocampal overexpression on this type of memory.

Overall, our results indicate that Aβ could be responsible for the decrease in Nr4a2 expression observed in AD mouse models and we provide direct proof supporting a key role for Nr4a2 in the oAβ-mediated synaptic dysfunction that leads to learning and memory impairments at early stages of AD, emerging as a potential therapeutic candidate for delaying the onset or progression for such an insidious pathology. These results support Nr4a2 as a putative target to be considered in future AD therapeutic approaches.

## Acknowledgments

We thank the UAB Servei d’Estabulari and Viral Vector Production (UPV) for technical support. This work was partially supported by grants from Ministerio de Ciencia e Innovación of Spain (PID2020-11751ORB-100), FEDER (Regional European Development Fund) and CIBERNED (CB06/05/0042) to J.R.A and Generalitat de Catalunya (SGR2021-0142) to C.S. P.E.C was supported by NIH grants R01 MH115543, R01MH125772, and R01 NS 113600. J.C.S was supported by a FPU fellowship from Ministerio de Economía y Competitividad, P.J.L. was supported by a Ruth L. Kirschstein National Research Service Award Fellowship F31MH10926, C.F was supported by a UAB-PIF fellowship.

